# Shifting effects of host physiological condition following pathogen establishment

**DOI:** 10.1101/2022.12.01.518758

**Authors:** Kate E. Langwig, A. Marm Kilpatrick, Macy J. Kailing, Nichole Laggan, J. Paul White, Heather M. Kaarakka, Jennifer A. Redell, John E. DePue, Katy L. Parise, Jeffrey T. Foster, Joseph R. Hoyt

## Abstract

Understanding host persistence with emerging pathogens is essential for conserving populations. Hosts may initially survive pathogen invasions through pre-adaptive mechanisms. However, whether pre-adaptive traits are directionally selected to increase in frequency depends on the heritability and environmental dependence of the trait and the costs of trait maintenance. Body condition is likely an important pre-adaptive mechanism aiding in host survival, although can be seasonally variable in wildlife hosts. We used data collected over seven years on bat body mass, infection, and survival to determine the role of host body condition during the invasion and establishment of the emerging disease, white-nose syndrome. We found that when the pathogen first invaded, bats with higher body mass were more likely to survive, but this effect dissipated following the initial epizootic. We also found that heavier bats lost more weight overwinter, but fat budgeting depended on infection severity. Lastly, we found little support that bat mass increased in the population after pathogen arrival, and there was high annual plasticity in individual bat masses. Overall, our results suggest that factors that contribute to host survival during pathogen invasion may diminish over time, and are potentially replaced by other host adaptations.

## Introduction

The introduction of novel pathogens to naive hosts can have profound effects on populations [1-7]. Hosts may survive initial pathogen invasion through multiple mechanisms including evading infection or pre-adaptive traits that allow for survival despite infection or disease [8, 9]. Importantly, factors enabling hosts to survive during initial invasion may not confer any advantage subsequently, particularly if pre-adaptive traits have strong tradeoffs, or are highly plastic (e.g. environmentally dependent) [10, 11]. Ultimately, traits that determine long-term host-pathogen coexistence may take longer to evolve and become widespread than traits allowing for initial survival, particularly if such traits provide stronger protection than preadaptive mechanisms [10, 12-14].

Factors that affect the probability of host survival with invasive pathogens include age, chronic disease, prior exposure, and body mass [15]. In general, hosts with adequate fat stores, high nutrient levels, and access to high quality habitat should demonstrate improved disease outcomes over weaker hosts. However, host body condition can be highly variable across seasons and years, even within individuals, leading to heterogeneity in the relationship between host body condition and disease, and making it a less reliable mechanism long-term [16]. Variable effects of body condition may be particularly pronounced when there is highly seasonal availability of food sources, leading to high stochasticity among individuals in their ability to consistently maintain high body condition when faced with annual disease outbreaks.

White-nose syndrome (WNS) is a seasonal annual epizootic of bats caused by the fungal pathogen *Pseudogymnoascus destructans* [17-20]. White-nose syndrome was first detected New York, USA in 2006, and has caused widespread declines in hibernating bat populations across North America [6, 21, 22]. *Pseudogymnoascus destructans* grows optimally in cool conditions (1–17 ºC) [23], resulting in annual winter epidemics that occur when bats begin hibernating [18]. Invasion of *P. destructans* into bat skin tissue causes severe physiological disruption, elevating bat metabolic rate and increasing respiratory acidosis [24, 25]. Bats, in turn, arouse to normalize blood pH which further increases evaporative water loss and causes dehydration. Higher energy expenditure from infection, increases fat loss, and emaciation, which frequently leads to mortality [26-28].

Increases in stored fat and improved budgeting of fat overwinter are therefore hypothesized to be important mechanisms determining bat survival with WNS [29-31], which typically increases within 4-5 years of WNS arrival after initially severe declines [6, 17, 32, 33]. However, other mechanisms of host persistence have also been described, including potential increases in host resistance through immunity or microbially-mediated reductions in pathogen growth [17, 33], and movement toward colder roosting conditions which limits fungal growth [34, 35]. Nonetheless, changes in body mass have the potential to have strong effects on bat survival, but comprehensive analyses on the effect of body mass on individual bat survival with WNS in the field have yet to be conducted. In addition, because host body condition may exhibit high annual variability [36], the importance of body mass as a sustained factor affecting population persistence with WNS merits additional investigation. Here, we investigate changes in the effect of body mass on survival of individual little brown bats (*Myotis lucifugus*) during the invasion and establishment of *P. destructans* across 24 sites. We hypothesized that while fatter bats might initially exhibit higher survival, the positive effects of higher body condition could diminish over time as host disease resistance increases in bat populations.

## Methods

We studied the arrival and establishment of *P. destructans* at 24 hibernacula (caves and mines where bats spend the winter) in Virginia, Wisconsin, Illinois, and Michigan over seven years (Tables S1-S3). We visited sites twice per winter and collected data on infection status and body mass of bats. At each site, we sampled up to 25 individual bats stratified across site sections. Because sites used in this study were primarily small mines where it was possible to observe all bats present, in many instances, all individuals in the population were sampled. For each bat, we collected a standardized epidermal swab sample [18], attached a unique aluminum band, and measured body mass using a digital scale (GDealer, accuracy +/-0.03 grams). Because common condition indices are no more effective than body mass for estimating fat stores [37], we did not include information on bat forearm size in order to reduce handling disturbance. At every visit, we recorded and resampled any previously banded bats present. We stored swabs in RNAlater until processing. We tested samples for *P. destructans* DNA using real-time PCR and quantified fungal loads [21, 38]. Animal handling protocols were approved by Virginia Tech IACUC (#17-180, #20-150).

We investigated the effect of bat early hibernation (November) body mass on the probability an individual was recaptured overwinter using a generalized linear mixed model (GLMM) with a binomial distribution and a probit link, with site as a random effect, and body mass and disease phase (epidemic = 1-3 years since pathogen arrival, or established = 4-7 years since pathogen arrival) as interacting fixed effects. Phases were established based on previous results demonstrating that populations approach stability by year 4 following WNS arrival [6, 32] For analyses of individual survival and body mass, results were similar whether we used categorical disease phase or years since WNS as a continuous variable (Appendix) and grouping by phase maximized the number of bats in the epidemic years when mortality was high and the number of recaptured bats was low. For bats that were recaptured overwinter, we examined the effect of early winter body mass and infection on the amount of mass lost overwinter during both the epidemic and established phase using a linear mixed model with site as a random effect and the change in body mass as the response variable and fixed effects of early winter mass interacting early winter fungal loads with additional additive effect of disease phase. Finally, we explored changes in mass over time since the invasion of *P. destructans* on an individual and population level to examine both plasticity and phenotypic change. For bats that were recaptured in multiple years, we used a linear mixed model with mass as a response variable, years since WNS as a fixed effect, and bat band ID as a random effect to explore plasticity in whether individual bat mass changed over time. At a population level, declines in sites with the best invasion mass data limited our ability to explore changes in mass, so we restricted our analyses to N=5 sites that were measured during invasion and had sufficient bats to estimate during established periods using log_10_ mass as our response variable (logged to normalize) and years since WNS interacting with season with site as a random effect.

## Results

As WNS invaded and caused massive declines in bat populations, bats that were heavier in early winter were more likely to be recaptured than lighter ones (Fig. 1; slope of mass ± SE: 0.320 ± 0.14, P = 0.0220). However, after WNS established in sites (years 4-7 following *P. destructans* detection), recapture overall was higher than during the epidemic (invasion vs establishment coef: 3.551 ± 1.46, P = 0.0152), and the effect of mass on the probability of recapture was significantly lower than the epidemic phase (interaction slope: -0.357 ± 0.16, P = 0.0250), and the slope did not differ significantly from 0 (Appendix 1.0.3).

**Figure 1.**
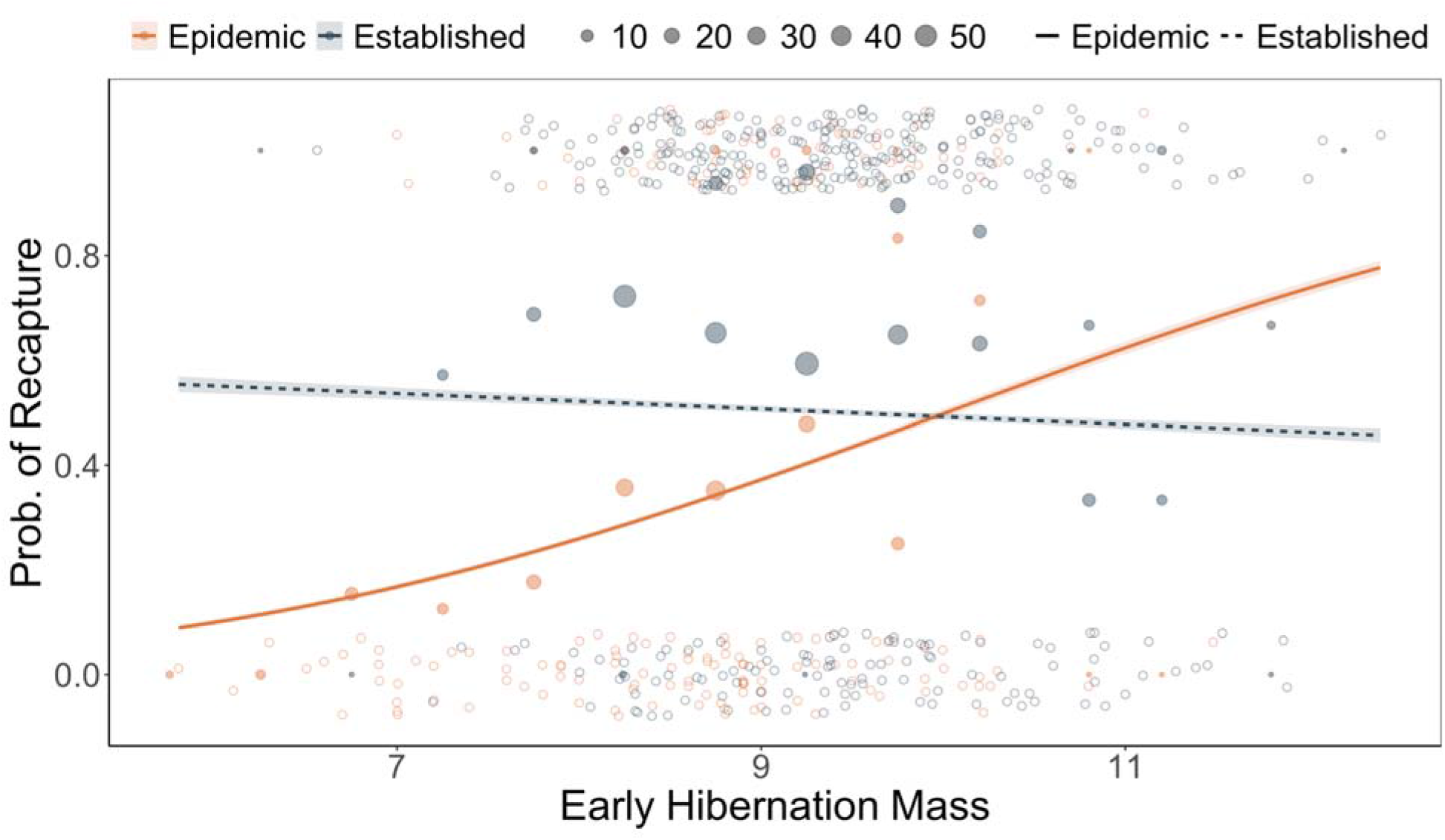
The effects of body mass during early hibernation on the probability of little brown bat recapture vary with time since *P. destructans* arrival. In years 0–3 post *P. destructans* arrival, the probability a bat was recaptured overwinter increased as early hibernation mass increased. However, after WNS established (>3 years since *P. destructans* arrival), there was no longer a clear trend between early hibernation body mass and bat survival. Solid points of early hibernation body masses during each phase show the fraction recaptured at 0.5 g bins (e.g. 9.75-10.25) and sample sizes for binned data.

For bats that survived overwinter and were recaptured, mass lost overwinter depended on both early hibernation weight and infection, and their interaction (Fig. 2A; P = 0.00948). There was little support for including disease phase as a predictor (P = 0.27), likely due to the paucity of bats recaptured during the epidemic phase when mortality was high (Table S2). Generally, bats that were heavier lost more weight overwinter than bats that were lighter (coef: -0.737 ± 0.16, t = -4.613). In addition, as infection increased, so did the amount of mass lost (coef: 1.002 ± 0.41, t = 2.467), but only for bats that were heavier in early winter; lighter bats lost less weight and weight loss did not vary with higher fungal loads (early mass:early loads coef: -0.106 +-0.04, t=-2.594, Appendix 2.0.2).

**Figure 2.**
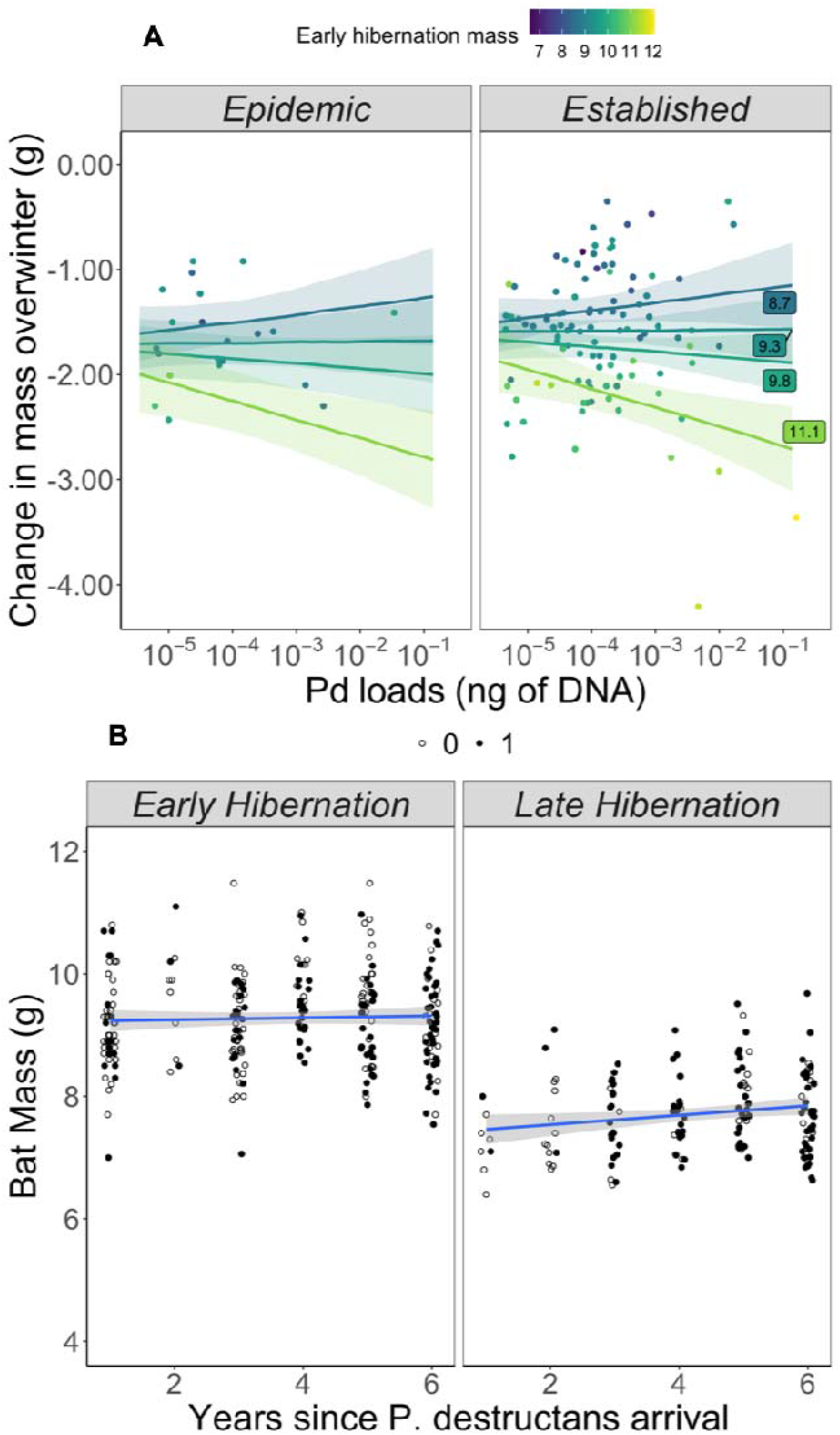
**(A)** Fungal loads and early hibernation (November) body mass of little brown bats strongly influences the change in individual bat mass over winter. Points show individual bats captured in both early and late hibernation. Colors denote masses of bats during early hibernation and labeled lines show predictions based on the 25th (8.7 g), 50th (9.3 g), 85th (9.8 g), and 95th (11.1 g) percentiles of the early hibernation masses. Bats that have higher initial body mass lose more weight over winter than bats with lower body mass (i.e. darker lines are higher), suggesting that bats budget fat stores accordingly over winter. In addition, fungal loads significantly modify the effect of early hibernation mass on mass lost overwinter. Bats with high infections that were heavier lose more mass than similarly infected bats that were lighter, suggesting that highly infected bats that survive to be recaptured budget fat in accordance with their infection status. **(B)** Average body mass of banded little brown bats in early (November) and late (March) that were recaptured (filled circles) or not recaptured (open circles) overwinter during the WNS epidemic (Years 0-3) and WNS established period (Years 4+) at the same sites over time (N=5). We found no clear support that hibernation body masses of bats increased over time when examining these data continuously (top) but marginal support categorically (Fig S1).

We found limited support for increases in mass at a population-level. Including years since pathogen arrival as a continuous effect, we found no clear support for increases in mass at a population-level (years since pathogen invasion coef: 0.002 ± 0.002, t=0.983, Fig. 2B, Table S3, Appendix 3.0.2). We did find support for a modest increase in log_10_ early hibernation body mass between the epidemic and established periods at 5 sites that were sampled at all time points in most years (established coef: 0.011 ± 0.005, t=2.031, Fig. S1, Appendix 3.0.4), however this was largely due to an increase between one annual time step (Year 3 to Year 4). We found no support for an increase in mass due to plasticity (Appendix 4). Using just recaptured bats, we found weak and unclear support for increases in log_10_ early hibernation body mass with years since WNS establishment (0.0037 ± 0.003, t=1.508, Fig. S1 closed circles, Table S2, Appendix 4.0.2). Furthermore, masses of individual bats that were recaptured in multiple years decreased non-significantly (−0.0023 ± 0.003, t=-0.701, Fig. S2, Appendix 4.03). Among individual bats recaptured annually, there was high plasticity in body mass which ranged from -1.78 : +1.09 g, suggesting that bat fat stores may be highly dependent on local conditions in summer and autumn.

## Discussion

We found that the effect of body mass on survival waned as the epidemic progressed. Furthermore, fat loss in bats increased with initial stored fat, as has been previously found in another species [39], suggesting that bats surviving with disease are budgeting fat stores to mitigate the physiological disruption posed by WNS. Importantly, we did not find evidence that bat survival once the disease established was enhanced by increases in the amount of stored fat [29]. We also found little support that fat increased at the population level as the disease established. When treating years since pathogen arrival continuously, there was no clear trend of increases in fat at the population-level. In some years, annual increases in fat occurred, but these were modest relative to the range of body conditions at the start of hibernation (recaptured bats during the established WNS period ranged from 7-12 grams and gains were an average of 0.18 grams. We also found no support of consistent mass increases in individual bats, and year to year fat stores were highly variable (range -1.78: +1.09 grams).

There are several potential reasons that could explain why the importance of fat changed as *P. destructans* established. First, the initial epizootic may have selected for fatter individuals, thus making the effects of fat less apparent as the pathogen established. However, body mass differences between the invasion and established phases were very modest relative to annual plasticity in bat masses, suggesting that this is unlikely. Second, bats in some populations have evolved higher pathogen resistance [33, 35] which may have reduced selection for increased body mass, particularly if fatter bats face other tradeoffs, such as reduced flight abilities [40, 41]. Third, bats have shifted to using cooler microclimates that also reduce the growth of the fungus, resulting in less severe disease [34]. Fourth, changes in the pathogen (e.g. a reduction in virulence) could have enabled more hosts to survive, thus experiencing fewer adverse effects (e.g. excess fat loss) from the pathogen [42, 43]. Lastly, bats may have adapted to the physiological disruption posed by infection, as evidenced by the relationship between mass loss, infection, and early hibernation weight. This finding is consistent with the hibernation optimization hypothesis [44-46], suggesting that bats do not use a fixed amount of fat during hibernation [47, 48], and generally aligns with findings conducted on unaffected little brown bats that demonstrated increases in arousals with increases in early hibernation fat [44]. Overall, increased fat stores may have been beneficial initially, but changes in other host or pathogen traits may have relaxed selection on fat over time.

Our results have important implications for the conservation of bats impacted by WNS. Supplemental feeding and enhancement of autumn bat habitat to increase insect prey abundance have been explored as a management strategy to increases bat fat stores to reduce WNS impacts [49]. Our results strongly suggest that while this may have been effective prior to or during pathogen invasion, it provides little benefit to bats once the pathogen has been established for several years. We find that bats budget fat in accordance with their infection severity and initial fat stores, suggesting that supplemental feeding might not achieve the desired benefit of enhancing bat survival if bats simply alter fat use accordingly. In addition, supplemental feeding of wildlife may have unexpected negative consequences, including increases in predation, increases in susceptibility due to less nutritious food sources, and enhancement of pathogen spread due to host aggregation [50], and these potential negative effects should be carefully considered before widescale implementation.

Species survival in the face of global change will likely require rapid adaptation and change itself may outpace the speed at which species can evolve [51, 52]. For species and populations that persist, some traits that may be beneficial for initial survival may prove less important over time [9, 53]. This phenomenon may be partly explained by coevolutionary theory which suggests that both hosts and pathogen must constantly adapt and innovate in order to maintain high fitness [12]. Ultimately, developing a more comprehensive understanding of the pre-adaptive factors that aid in population health can enable us to build more resilient populations in the Anthropocene.

## Acknowledgements

We thank Steffany Yamada for data curation support, Rick Reynolds for logistical support, and the many landowners for site access.

## Data Availability Statement

The datasets and code generated in this study have been included in the electronic supplementary material for review and will be deposited in Dryad Digital Repository upon final submission. Exact site locations are not disclosed to protect endangered species and landowners.

## Ethics Statement

Animal handling protocols were approved by Virginia Tech IACUC (#17-180, #20-150). Field work was conducted under approved permits from the Wisconsin Department of Natural Resources, the Virginia Division of Game and Inland Fisheries, the Illinois Department of Natural Resources, and the Michigan Department of Natural Resources. All personnel followed field hygiene protocols for *P. destructans* as recommended by the USFWS.

## Author Contributions

K.E.L.: conceptualization, investigation, methodology, funding acquisition, resources, project administration, data curation, formal analysis, writing-original draft, writing-review and editing;

M.J.K.: investigation, methodology, writing-review and editing,

N.A.L.: investigation, methodology, writing-review and editing,

J.P.W.: investigation, writing-review and editing,

H.M.K.: investigation, writing-review and editing,

J.A.R.: investigation, writing-review and editing,

J.E.D.: investigation, resources

K.L.P.: investigation, writing-review and editing,

J.T.F.: investigation, funding acquisition, writing-review and editing,

A.M.K.: investigation, funding acquisition, writing-review and editing,

J.R.H.: conceptualization, investigation, methodology, funding acquisition, resources, project administration, data curation, writing-review and editing

