## Supplement for "Shifting effects of host physiological condition following pathogen establishment"

Langwig

Supplemental Material

**See Appendix for Statistical Output.**

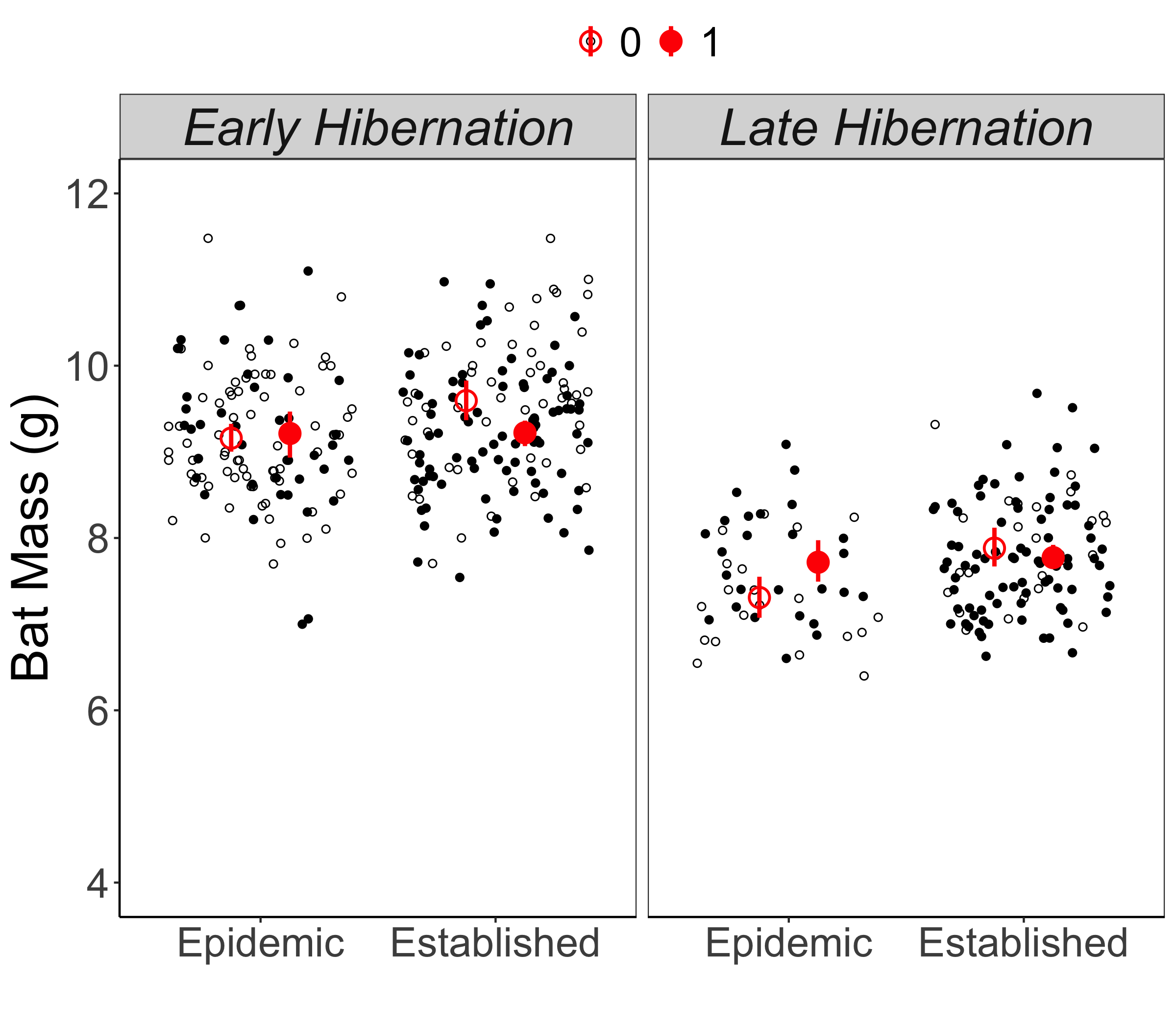

Figure Supplement 1. Average body mass of banded little brown bats in early (November) and late (March) that were recaptured (filled circles) or not recaptured (open circles) overwinter during the WNS epidemic (Years 0-3) and WNS established period (Years 4+) at the same sites over time (N=5).

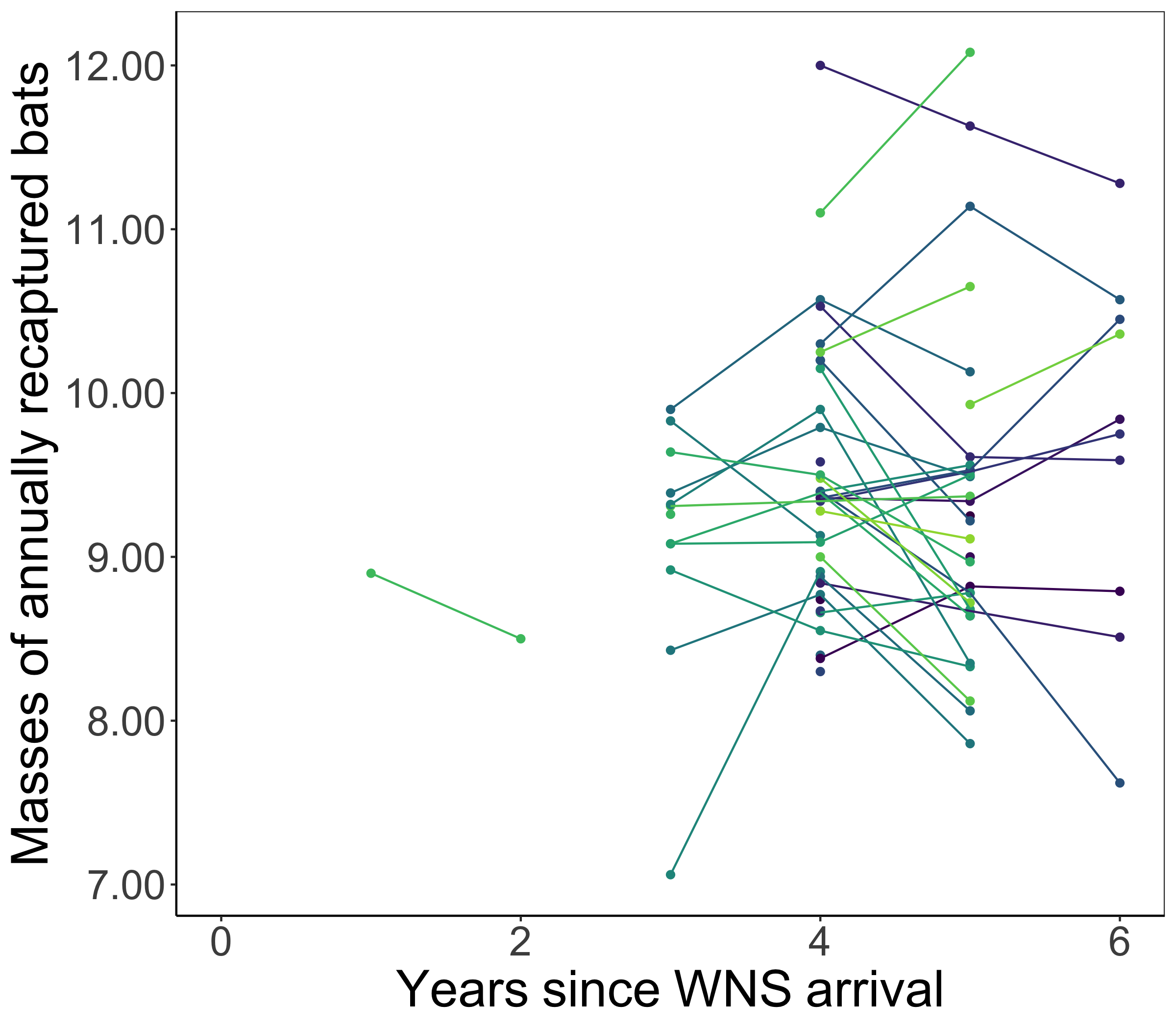

Figure Supplement 2. Body mass of individual little brown bats that were recaptured in at least two years in early (November) hibernation. Colors show individual bats and points display bat mass at recapture and lines link between recapture events over time. We found no support that early hibernation body masses of bats changed over time but individual variation in weights annually was high (range -1.78 : +1.09 g).

Table S1. Sample sizes of bats for analysis shown in Fig. 1.

| state | site | years since pathogen arrival | N |
| --- | --- | --- | --- |
| MI | ATENS | 0 | 3 |
| MI | ATENS | 1 | 1 |
| MI | ATENS | 3 | 14 |
| MI | ATENS | 4 | 14 |
| MI | ATENS | 5 | 14 |
| MI | ATENS | 6 | 7 |
| MI | ATENS | 7 | 8 |
| MI | EN AD | 0 | 5 |
| MI | EN AD | 1 | 14 |
| MI | EN AD | 2 | 2 |
| MI | EN AD | 3 | 10 |
| MI | EN AD | 4 | 20 |
| MI | EN AD | 5 | 1 |
| MI | EN AD | 6 | 1 |
| MI | EN AD | 7 | 1 |
| MI | SS C | 0 | 3 |
| MI | SS C | 1 | 1 |
| MI | SS C | 3 | 18 |
| MI | SS C | 4 | 12 |
| MI | SS C | 5 | 10 |
| MI | SS C | 6 | 10 |
| MI | SS C | 7 | 7 |
| MI | UTH L | 5 | 20 |
| MI | UTH L | 6 | 24 |
| MI | UTH L | 7 | 50 |
| MI | YLOR | 0 | 3 |
| MI | YLOR | 1 | 3 |
| MI | YLOR | 2 | 23 |
| MI | YLOR | 3 | 23 |
| MI | YLOR | 4 | 23 |
| MI | YLOR | 5 | 20 |
| MI | YLOR | 6 | 28 |
| VA | ARR C | 2 | 14 |
| VA | EATHI | 3 | 1 |
| VA | LLY | 0 | 10 |
| VA | ONELE | 2 | 1 |
| VA | OSSRO | 1 | 1 |
| VA | PMANS | 2 | 11 |
| VA | WBERR | 3 | 10 |
| VA | WNEYS | 2 | 6 |
| WI | . JOH | 0 | 2 |
| WI | . JOH | 2 | 5 |
| WI | AR CR | 5 | 2 |
| WI | AR CR | 6 | 3 |
| WI | DGEVI | 2 | 1 |
| WI | DGEVI | 3 | 3 |
| WI | DGEVI | 4 | 1 |
| WI | DGEVI | 5 | 1 |
| WI | IGER | 1 | 20 |
| WI | IGER | 2 | 13 |
| WI | IGER | 3 | 4 |
| WI | IGER | 4 | 2 |
| WI | IGER | 5 | 3 |
| WI | MP ST | 1 | 2 |
| WI | MP ST | 3 | 2 |
| WI | RIBEL | 0 | 5 |
| WI | RIBEL | 1 | 3 |
| WI | RIBEL | 2 | 4 |
| WI | RIBEL | 3 | 24 |
| WI | RIBEL | 4 | 22 |
| WI | RIBEL | 5 | 19 |
| WI | RIBEL | 6 | 15 |
| WI | ROY S | 0 | 1 |
| WI | ROY S | 1 | 3 |
| WI | ROY S | 2 | 1 |
| WI | ROY S | 3 | 19 |
| WI | ROY S | 4 | 20 |
| WI | ROY S | 5 | 20 |
| WI | ROY S | 6 | 21 |
| WI | RSESH | 1 | 1 |
| WI | RSESH | 3 | 2 |
| WI | RSESH | 4 | 2 |
| WI | RSESH | 5 | 1 |
| WI | RSESH | 7 | 17 |
| WI | SCOBE | 3 | 7 |
| WI | SCOBE | 4 | 3 |
| WI | SCOBE | 5 | 4 |
| WI | SCOBE | 6 | 1 |
| WI | SCOBE | 7 | 3 |
| WI | ST RI | 1 | 1 |
| WI | UTH P | 1 | 32 |
| WI | UTH P | 2 | 8 |
| WI | UTH P | 3 | 2 |
| WI | UTH P | 5 | 3 |

Table S2. Sample sizes of recaptured bats used in analysis in Fig. 2A.

| state | site | years since pathogen arrival | N |
| --- | --- | --- | --- |
| MI | ATENS | 4 | 8 |
| MI | ATENS | 5 | 10 |
| MI | ATENS | 6 | 5 |
| MI | EN AD | 4 | 15 |
| MI | EN AD | 5 | 1 |
| MI | SS C | 4 | 10 |
| MI | SS C | 5 | 7 |
| MI | SS C | 6 | 6 |
| MI | UTH L | 5 | 3 |
| MI | UTH L | 6 | 7 |
| MI | YLOR | 3 | 14 |
| MI | YLOR | 4 | 20 |
| MI | YLOR | 5 | 16 |
| VA | LLY | 0 | 2 |
| VA | PMANS | 2 | 4 |
| WI | DA MI | 6 | 1 |
| WI | IDEN | 5 | 2 |
| WI | IGER | 1 | 2 |
| WI | IGER | 2 | 1 |
| WI | RIBEL | 4 | 3 |
| WI | RIBEL | 5 | 3 |
| WI | ROY S | 3 | 5 |
| WI | ROY S | 5 | 4 |
| WI | RSESH | 4 | 1 |
| WI | SCOBE | 4 | 1 |
| WI | UTH P | 2 | 2 |
| WI | Y CIT | 5 | 5 |

Table S3. Sample sizes for analysis used in Figure 2.

| state | site | years since pathogen arrival | N | mean mass |
| --- | --- | --- | --- | --- |
| MI | YLOR | 3 | 39 | 8.45 |
| MI | YLOR | 4 | 42 | 8.69 |
| MI | YLOR | 5 | 37 | 8.48 |
| MI | YLOR | 6 | 53 | 8.42 |
| WI | IGER | 1 | 22 | 9.1 |
| WI | IGER | 2 | 3 | 8.49 |
| WI | RIBEL | 3 | 11 | 9.44 |
| WI | RIBEL | 4 | 14 | 9.17 |
| WI | RIBEL | 5 | 19 | 9.43 |
| WI | RIBEL | 6 | 9 | 9.4 |
| WI | ROY S | 2 | 2 | 7.05 |
| WI | ROY S | 3 | 25 | 8.48 |
| WI | ROY S | 4 | 3 | 8.27 |
| WI | ROY S | 5 | 27 | 8.73 |
| WI | ROY S | 6 | 42 | 8.41 |
| WI | UTH P | 1 | 37 | 8.8 |
| WI | UTH P | 2 | 19 | 8.57 |
| WI | UTH P | 3 | 3 | 9.02 |
| WI | UTH P | 5 | 1 | 9.92 |
