## Appendix for "Shifting effects of host physiological condition following pathogen establishment": Appendix_RNotebook.html

R Notebook - Langwig WNS fat ms


Code 

- Show All Code
- Hide All Code
- Download Rmd

### R Notebook - Langwig WNS fat ms

- 1 Figure 1: Recapture analyses - Does body mass influence bat overwinter recapture? How does this change as the epizootic progresses?
  - 1.0.1 What data does this include? (See also table S1)
  - 1.0.2 Continuous version
  - 1.0.3 Categorical version - Anaysis for Figure 1
- 2 Figure 2A - Change in weight by early winter weight, loads, and years since WNS
  - 2.0.1 what’s the data? (See table S2)
  - 2.0.2 Figure 2A - does the change in weight change with ysw, fungal loads, and early weight?
  - 2.0.3 PLOT of Model 2A - there is effect of loads and starting weight on weight loss, what does the plot look like?
- 3 Fig 2B, Fig S1 - Fat over time - Do we find that bats are getting fatter over time?
  - 3.0.1 what’s the data? (See table S3)
  - 3.0.2 all banded bats plotted - epidemic & established, late & early winter
  - 3.0.3 Plotted model results and data - Figure 2B
  - 3.0.4 Categorical analysis of fat
- 4 Figure S2 - How much individual plasticity is there in bat weight?
  - 4.0.1 data for analysis - core midwest sampled across epidemic, but JUST recaptured bats (same as Figure 2A)
  - 4.0.2 is there an effect of years since WNS on early & late hiber mass?
  - 4.0.3 Figure S2 - annual plasticity in bat mass - For bats that were recaptured annually, how do their weights change?
  - 4.0.4 the plot doesn’t look like there is a change - what do the stats say?

### 1 Figure 1: Recapture analyses - Does body mass influence bat overwinter recapture? How does this change as the epizootic progresses?

##### 1.0.1 What data does this include? (See also table S1)


```
#data is all bats banded in 24 different sites that were surveyed in early and late hibernation
unique(str_sub(m.early$site, 3,7))
```


```
 [1] "PMANS" "EATHI" "OSSRO" "WNEYS" "ONELE" "LLY"   "ARR C"
 [8] "WBERR" ". JOH" "EN AD" "ATENS" "SS C"  "YLOR " "RIBEL"
[15] "RSESH" "ROY S" "MP ST" "ST RI" "IGER " "UTH P" "SCOBE"
[22] "DGEVI" "UTH L" "AR CR"
```

##### 1.0.2 Continuous version


```
mod9 = glm(EverRecapturedYN~mass*ysw,family=binomial(),data=m.early, subset=ysw>0);summary(mod9)
```


```
Call:
glm(formula = EverRecapturedYN ~ mass * ysw, family = binomial(), 
    data = m.early, subset = ysw > 0)

Deviance Residuals: 
    Min       1Q   Median       3Q      Max  
-1.8438  -1.2374   0.8605   0.9539   2.0274  

Coefficients:
            Estimate Std. Error z value Pr(>|z|)    
(Intercept) -9.46377    2.34487  -4.036 5.44e-05 ***
mass         0.99530    0.25687   3.875 0.000107 ***
ysw          1.84986    0.49476   3.739 0.000185 ***
mass:ysw    -0.18162    0.05319  -3.414 0.000640 ***
---
Signif. codes:  0 ‘***’ 0.001 ‘**’ 0.01 ‘*’ 0.05 ‘.’ 0.1 ‘ ’ 1

(Dispersion parameter for binomial family taken to be 1)

    Null deviance: 695.03  on 515  degrees of freedom
Residual deviance: 660.45  on 512  degrees of freedom
  (227 observations deleted due to missingness)
AIC: 668.45

Number of Fisher Scoring iterations: 4
```


```
library(effects)
plot(allEffects(mod9))
```


```
library(viridis)
mass_and_ysw_continous_plot=ggplot(m.early, aes(x=mass, y=EverRecapturedYN,color=ysw)) +
  geom_point(size=5,alpha=.5)+ 
  geom_line(data=subset(nd,ysw==1),aes(x=mass,y=fit,color=ysw),size=1)+#
  geom_line(data=subset(nd,ysw==3),aes(x=mass,y=fit,color=ysw),size=1)+#
  geom_line(data=subset(nd,ysw==5),aes(x=mass,y=fit,color=ysw),size=1)+#
  geom_line(data=subset(nd,ysw==7),aes(x=mass,y=fit,color=ysw),size=1)+#
  geom_line(data=subset(nd,ysw==9),aes(x=mass,y=fit,color=ysw),size=1)+#
  ylab("Prob. of Recapture")+
  xlab(expression(Early~Hibernation~Mass))+
  scale_colour_viridis(option="inferno")+
  theme_bw()+
  theme(panel.grid = element_blank(),axis.title=element_text(size=25),
        axis.text=element_text(size=20), axis.line=element_line(),
        legend.position="right",legend.text = element_text(size=20,face="italic"),legend.title = element_blank(),strip.text = element_text(size=25,face="italic"))

mass_and_ysw_continous_plot
```


```
Warning: Removed 249 rows containing missing values (geom_point).
```

##### 1.0.3 Categorical version - Anaysis for Figure 1


```
mod10a = glmer(EverRecapturedYN~mass*ysw.cat + (1|site),
              family=binomial(link="probit"),
              control=glmerControl(optimizer="bobyqa", optCtrl=list(maxfun=100000)),
                                   data=m.early)
summary(mod10a)
```


```
Generalized linear mixed model fit by maximum likelihood
  (Laplace Approximation) [glmerMod]
 Family: binomial  ( probit )
Formula: EverRecapturedYN ~ mass * ysw.cat + (1 | site)
   Data: m.early
Control: 
glmerControl(optimizer = "bobyqa", optCtrl = list(maxfun = 1e+05))

     AIC      BIC   logLik deviance df.resid 
   599.6    620.9   -294.8    589.6      521 

Scaled residuals: 
    Min      1Q  Median      3Q     Max 
-3.0904 -0.6028  0.3341  0.6738  3.1400 

Random effects:
 Groups Name        Variance Std.Dev.
 site   (Intercept) 0.5681   0.7538  
Number of obs: 526, groups:  site, 22

Fixed effects:
                        Estimate Std. Error z value Pr(>|z|)   
(Intercept)              -3.2035     1.2417  -2.580  0.00988 **
mass                      0.3198     0.1396   2.291  0.02198 * 
ysw.catEstablished        3.5513     1.4625   2.428  0.01517 * 
mass:ysw.catEstablished  -0.3566     0.1586  -2.248  0.02459 * 
---
Signif. codes:  0 ‘***’ 0.001 ‘**’ 0.01 ‘*’ 0.05 ‘.’ 0.1 ‘ ’ 1

Correlation of Fixed Effects:
            (Intr) mass   ysw.cE
mass        -0.983              
ysw.ctEstbl -0.847  0.851       
mss:ysw.ctE  0.853 -0.868 -0.989
```


###### 1.0.3.1 standard errors calcs for plotting


```
#use bootMer to get std errors
mySumm <- function(.) {
  predict(., newdata=nd2, type="response",re.form=NA)
}

boot10a <- lme4::bootMer(mod10a, mySumm, nsim=250, use.u=TRUE, type="parametric")
std.err <- apply(boot10a$t, 2, std.error)
```


```
# prediction model
nd2$yhat = predict(mod10a, newdata=nd2, type="response", re.form=NA)
nd2 = cbind(nd2,std.err)
```

###### 1.0.3.2 full plot- Figure 1


```
mass_and_ysw_categorical_plot=ggplot(m.early.no.pre, aes(x=mass, y=EverRecapturedYN,fill=ysw.cat)) +
   geom_jitter(data=m.early.no.pre, aes(color=ysw.cat),height = .08, size=2,alpha=.5,shape=1)+ 
  geom_point(data=mass.summary.fine, aes(x=mass.round, y=EverRecapturedYN,color=ysw.cat,size=N.bats.in.fat.cat),alpha=0.5)+
  geom_line(data=nd2, aes(x=mass, y=EverRecapturedYN,color=ysw.cat, linetype=ysw.cat),size=1)+
  geom_ribbon(data=nd2, aes(x=mass, ymin = EverRecapturedYN-(std.err*1.96), ymax = EverRecapturedYN+(std.err*1.96)), alpha=0.2) +
  ylab("Prob. of Recapture")+
  xlab(expression(Early~Hibernation~Mass))+
  colScale+
  fillScale+
  theme_bw()+
  theme(panel.grid = element_blank(),axis.title=element_text(size=25),
        axis.text=element_text(size=20), axis.line=element_line(),
        legend.position="top",
        legend.text = element_text(size=20),#,face="italic"
        legend.title = element_blank(),
        strip.text = element_text(size=25,face="italic"))

mass_and_ysw_categorical_plot
```


```
Warning: Removed 249 rows containing missing values (geom_point).
```

### 2 Figure 2A - Change in weight by early winter weight, loads, and years since WNS

##### 2.0.1 what’s the data? (See table S2)


```
m2 = m %>% 
  group_by(wyear, band)%>%
  filter(n()>1) %>%
  arrange(pdate) %>%
  mutate(dloads = gdL - lag(gdL, default = NA), 
         dweight = mass - lag(mass, default=NA),
         early.weight = lag(mass, default=NA),
         early.gdL = lag(gdL, default=NA),
         early.temp = lag(temp, default=NA))
m2$load.change = m2$dloads+(min(m2$dloads,na.rm=T)+1)
m2$ysw.f = as.factor(m2$ysw)
m2$ysw.cat[m2$ysw>-1& m2$ysw<3] = "Epidemic"
```


```
Warning: Unknown or uninitialised column: `ysw.cat`.
```


```
m2$ysw.cat[m2$ysw>2] = "Established" 
 
m2e = m2 %>%
  group_by(band,wyear)%>%
  arrange(pdate)%>%
  mutate(change.late.loads = lead(load.change,1))
m2e = subset(m2e, season == "hiber_earl")
#m2e did bring in the load change data

m2l = subset(m2, season=="hiber_late")
##m2l should have a bunch of columns matched in
#remove bats that got 'fatter' over the season?
#only 10; mostly from wet sites
#so probably moisture
unique(m2l$dweight)
```


```
  [1]    NA -0.18  0.20 -1.37 -1.16 -2.47 -1.82 -2.01 -0.95 -1.64
 [11] -1.41 -2.09 -1.54 -1.57 -2.35 -0.96 -0.68 -1.09 -1.69 -0.57
 [21] -0.93 -1.29 -1.43 -1.38 -0.87 -0.47 -0.78 -0.50 -0.15 -1.72
 [31] -0.99 -2.34 -1.93 -2.10 -2.26 -1.75 -1.76 -2.71 -2.05 -2.42
 [41] -1.61 -0.92 -1.75 -1.59 -2.12 -2.18 -1.59 -2.92 -2.23 -2.78
 [51] -4.21 -1.81 -1.61 -1.55 -2.06 -0.72 -1.58 -1.83 -1.87 -2.24
 [61] -1.55 -1.58 -1.44 -1.80 -2.66 -0.85 -1.18 -1.48 -1.40 -2.72
 [71] -0.91 -1.47 -2.30  0.10 -1.42 -2.27 -1.26 -1.25 -0.60 -1.45
 [81] -1.14 -3.36 -2.11 -1.54 -2.08 -2.41 -1.52 -1.96 -1.72 -1.21
 [91] -1.06 -1.65 -2.79 -1.50 -1.23 -1.80 -1.68 -1.85 -1.03 -1.91
[101] -1.19 -1.24 -1.87 -2.43  0.26 -1.36  0.76 -2.45 -1.45 -1.30
[111] -1.09 -0.95 -2.11 -1.73 -1.41 -1.07 -1.33 -1.20 -0.35 -1.21
[121] -1.42 -2.64 -1.73 -1.53 -0.77 -0.80 -1.02 -1.89 -0.83 -2.02
[131] -1.99 -2.07 -2.10  0.00 -1.80
```


```
m2l = m2l %>%
  filter(dweight<0.1)

#final dataset is all banded bats (all states) recaptured, filtered for obvious mistakes (bats that gained weight, mostly from very wet sites) with epidemic as years 0-3, and established as 4 and on
```

##### 2.0.2 Figure 2A - does the change in weight change with ysw, fungal loads, and early weight?


```
w5 = lmer(dweight~early.weight*early.lgdL+ early.weight+ysw.cat+(1|site), data=m2l)
summary(w5)
```


```
Linear mixed model fit by REML ['lmerMod']
Formula: 
dweight ~ early.weight * early.lgdL + early.weight + ysw.cat +  
    (1 | site)
   Data: m2l

REML criterion at convergence: 207.1

Scaled residuals: 
     Min       1Q   Median       3Q      Max 
-2.91255 -0.63025 -0.07698  0.59459  2.93841 

Random effects:
 Groups   Name        Variance Std.Dev.
 site     (Intercept) 0.01191  0.1091  
 Residual             0.23989  0.4898  
Number of obs: 134, groups:  site, 17

Fixed effects:
                        Estimate Std. Error t value
(Intercept)              5.19922    1.58361   3.283
early.weight            -0.76018    0.15904  -4.780
early.lgdL               1.06262    0.40360   2.633
ysw.catEstablished       0.39573    0.20396   1.940
early.weight:early.lgdL -0.11196    0.04083  -2.742

Correlation of Fixed Effects:
            (Intr) erly.w erly.L ysw.cE
early.weght -0.987                     
early.lgdL   0.952 -0.950              
ysw.ctEstbl -0.076 -0.040  0.029       
erly.wgh:.L -0.943  0.953 -0.994 -0.026
```


```
plot(allEffects(w5))
```

##### 2.0.3 PLOT of Model 2A - there is effect of loads and starting weight on weight loss, what does the plot look like?


```
mass_over_winter_plot=ggplot(m2l, aes(x=early.lgdL, y=dweight, fill=early.weight, group=early.weight)) +
  facet_wrap(~ysw.cat)+
  geom_point(aes(color=early.weight))+
    geom_line(data=newdat2, aes(x=early.lgdL, y=dweight,color=early.weight, fill=early.weight),size=1)+#linetype="dashed"
  geom_ribbon(data=newdat2, aes(x=early.lgdL, ymin = dweight-(std.err*1.96), ymax = dweight+(std.err*1.96)), alpha=0.2 ) +
  ylab("Mass loss overwinter (g)")+
  xlab(expression(Pd~loads~"(ng of DNA)"))+
  geom_label_repel(data=subset(newdat2, ysw.cat=="Established"), aes(label=label))+
  scale_y_continuous(labels = scales::number_format(accuracy = 0.01))+
  scale_x_continuous(limits=c(-5.5,-0.5),breaks=c(-5,-4,-3,-2,-1), labels=expression(10^-5,10^-4,10^-3,10^-2, 10^-1))+
  scale_colour_viridis(option="D")+#, discrete=T
  scale_fill_viridis(option="D")+#, discrete=T
  labs(colour="Early hibernation mass", fill="Early hibernation mass")+
  theme_bw()+
  theme(panel.grid = element_blank(),axis.title=element_text(size=25),
        axis.text=element_text(size=20), axis.line=element_line(),
        legend.position="top",
        legend.text = element_text(size=12),#,face="italic"
        legend.title = element_text(size=15),
       # legend.title = element_blank(),
        strip.text = element_text(size=25,face="italic"))
```


```
Warning: Ignoring unknown aesthetics: fill
```


```
mass_over_winter_plot
```


```
Warning: Removed 34 rows containing missing values (geom_point).
Warning: Removed 8 row(s) containing missing values (geom_path).
Warning: Removed 192 rows containing missing values (geom_label_repel).
```

### 3 Fig 2B, Fig S1 - Fat over time - Do we find that bats are getting fatter over time?

##### 3.0.1 what’s the data? (See table S3)


```
#this dataset has 5 sites with the best data of the  populations sampled throughout multiple epidemic years and spanning the categorical time periods
unique(str_sub(small.fat$site, 3,7))
```

##### 3.0.2 all banded bats plotted - epidemic & established, late & early winter


```
mod.test8b = lmer(log10(mass)~ysw*season + (1|site), data=small.fat); summary(mod.test8b)
```


```
Linear mixed model fit by REML ['lmerMod']
Formula: log10(mass) ~ ysw * season + (1 | site)
   Data: small.fat

REML criterion at convergence: -1524.1

Scaled residuals: 
    Min      1Q  Median      3Q     Max 
-3.3358 -0.6858 -0.0625  0.7584  2.7425 

Random effects:
 Groups   Name        Variance  Std.Dev.
 site     (Intercept) 0.0001007 0.01004 
 Residual             0.0012507 0.03536 
Number of obs: 408, groups:  site, 5

Fixed effects:
                      Estimate Std. Error t value
(Intercept)           0.961465   0.007573 126.953
ysw                   0.001623   0.001651   0.983
seasonhiber_late     -0.094977   0.010137  -9.370
ysw:seasonhiber_late  0.003798   0.002260   1.680

Correlation of Fixed Effects:
            (Intr) ysw    ssnhb_
ysw         -0.741              
seasnhbr_lt -0.381  0.414       
ysw:ssnhbr_  0.376 -0.483 -0.931
```


```
#continous version - no support
```

##### 3.0.3 Plotted model results and data - Figure 2B


```
fat.over.time.continuous.small = ggplot(data=small.fat, aes(x=ysw, y=mass))+
  facet_wrap(~season2)+
  #geom_point()+
  geom_jitter(aes(shape=as.factor(RecapturedSameYearYN)),width=0.1, height=0)+
  stat_smooth(method="lm")+
  xlab("Years since P. destructans arrival")+
  ylab("Bat Mass (g)")+
  ylim(4,12)+
#  stat_summary(fun.data = "mean_cl_boot",position = position_dodge(width = 0.5), colour = "red", size = 1)+
  scale_colour_viridis(option="D")+#, discrete=T
  scale_fill_viridis(option="D")+#, discrete=T
  scale_shape_manual(values=c(1,19))+
  theme_bw()+
  theme(panel.grid = element_blank(),axis.title=element_text(size=25),
        axis.text=element_text(size=20), axis.line=element_line(),
        legend.position="top",
        legend.text = element_text(size=20),#,face="italic"
        legend.title = element_blank(),
        strip.text = element_text(size=25,face="italic"))
fat.over.time.continuous.small
```


```
`geom_smooth()` using formula 'y ~ x'
```

##### 3.0.4 Categorical analysis of fat


```
mod.test8b = lmer(log10(mass)~ysw.cat*season + (1|site), data=small.fat); summary(mod.test8b)
```


```
Linear mixed model fit by REML ['lmerMod']
Formula: log10(mass) ~ ysw.cat * season + (1 | site)
   Data: small.fat

REML criterion at convergence: -1530.8

Scaled residuals: 
    Min      1Q  Median      3Q     Max 
-3.3183 -0.6780 -0.0357  0.7383  2.8325 

Random effects:
 Groups   Name        Variance  Std.Dev.
 site     (Intercept) 8.399e-05 0.009165
 Residual             1.246e-03 0.035299
Number of obs: 408, groups:  site, 5

Fixed effects:
                                     Estimate Std. Error t value
(Intercept)                          0.961836   0.005341 180.074
ysw.catEstablished                   0.010447   0.005143   2.031
seasonhiber_late                    -0.083343   0.006210 -13.421
ysw.catEstablished:seasonhiber_late  0.006032   0.007655   0.788

Correlation of Fixed Effects:
            (Intr) ysw.cE ssnhb_
ysw.ctEstbl -0.462              
seasnhbr_lt -0.325  0.384       
ysw.ctEst:_  0.268 -0.540 -0.804
```


```
#categorical version - some support but effect is small
```


```
fat.over.time.small.categorical = ggplot(data=small.fat, aes(x=ysw.cat, y=mass,shape=as.factor(RecapturedSameYearYN)))+
  facet_wrap(~season2)+
  geom_jitter()+
  xlab("")+
  ylab("Bat Mass (g)")+
  ylim(4,12)+
  stat_summary(fun.data = "mean_cl_boot",position = position_dodge(width = 0.5), colour = "red", size = 1)+
  scale_colour_viridis(option="D")+#, discrete=T
  scale_fill_viridis(option="D")+#, discrete=T
  scale_shape_manual(values=c(1,19))+
  theme_bw()+
  theme(panel.grid = element_blank(),axis.title=element_text(size=25),
        axis.text=element_text(size=20), axis.line=element_line(),
        legend.position="top",
        legend.text = element_text(size=20),#,face="italic"
        legend.title = element_blank(),
        strip.text = element_text(size=25,face="italic"))
fat.over.time.small.categorical
```

### 4 Figure S2 - How much individual plasticity is there in bat weight?

##### 4.0.1 data for analysis - core midwest sampled across epidemic, but JUST recaptured bats (same as Figure 2A)


```
m2.mw.fat = m2 %>%
  filter(state=="MI"|state=="WI"|state=="IL")%>%
  drop_na(mass)
```

##### 4.0.2 is there an effect of years since WNS on early & late hiber mass?


```
#analysis of just recaptured bats
#no effect of years since WNS
mod.test5 = lmer(log10(mass)~ysw*season + (1|site), data=m2.mw.fat)
summary(mod.test5)
```


```
Linear mixed model fit by REML ['lmerMod']
Formula: log10(mass) ~ ysw * season + (1 | site)
   Data: m2.mw.fat

REML criterion at convergence: -1127.9

Scaled residuals: 
    Min      1Q  Median      3Q     Max 
-3.5806 -0.6766  0.0152  0.6238  2.7049 

Random effects:
 Groups   Name        Variance  Std.Dev.
 site     (Intercept) 1.739e-05 0.00417 
 Residual             1.732e-03 0.04162 
Number of obs: 331, groups:  site, 14

Fixed effects:
                      Estimate Std. Error t value
(Intercept)           0.953711   0.010637  89.662
ysw                   0.003715   0.002464   1.508
seasonhiber_late     -0.067333   0.017911  -3.759
ysw:seasonhiber_late -0.002464   0.004029  -0.612

Correlation of Fixed Effects:
            (Intr) ysw    ssnhb_
ysw         -0.943              
seasnhbr_lt -0.548  0.530       
ysw:ssnhbr_  0.537 -0.575 -0.966
```

##### 4.0.3 Figure S2 - annual plasticity in bat mass - For bats that were recaptured annually, how do their weights change?


```
change.weight.individuals = ggplot(m2.annual, aes(x=ysw, y=mass,color=band))+
  geom_point()+
  geom_line()+
  xlab(expression(Years~since~WNS~arrival))+
  ylab("Masses of annually recaptured bats")+
  scale_y_continuous(labels = scales::number_format(accuracy = 0.01))+
  scale_x_continuous(limits=c(0,6))+
  scale_colour_viridis(option="D",discrete=T)+#, 
  scale_fill_viridis(option="D",discrete=T)+#, discrete=T
  theme_bw()+
  theme(panel.grid = element_blank(),axis.title=element_text(size=25),
        axis.text=element_text(size=20), axis.line=element_line(),
        legend.position="none",
        legend.text = element_text(size=12),#,face="italic"
        legend.title = element_text(size=15),
        # legend.title = element_blank(),
        strip.text = element_text(size=25,face="italic"))

change.weight.individuals
```


```
Warning: Removed 56 rows containing missing values (geom_point).
Warning: Removed 56 row(s) containing missing values (geom_path).
```

##### 4.0.4 the plot doesn’t look like there is a change - what do the stats say?


```
a1 = lmer(log10(mass)~ysw + (1|band), data = m2.annual)
summary(a1)
```


```
Linear mixed model fit by REML ['lmerMod']
Formula: log10(mass) ~ ysw + (1 | band)
   Data: m2.annual

REML criterion at convergence: -339.3

Scaled residuals: 
     Min       1Q   Median       3Q      Max 
-2.78653 -0.40482 -0.03267  0.53830  1.63213 

Random effects:
 Groups   Name        Variance  Std.Dev.
 band     (Intercept) 0.0011386 0.03374 
 Residual             0.0005504 0.02346 
Number of obs: 91, groups:  band, 42

Fixed effects:
             Estimate Std. Error t value
(Intercept)  0.977716   0.015468  63.210
ysw         -0.002313   0.003298  -0.701

Correlation of Fixed Effects:
    (Intr)
ysw -0.926
```
